# A topographic visual pathway into the central brain of Drosophila

**DOI:** 10.1101/183707

**Authors:** Lorin Timaeus, Laura Geid, Thomas Hummel

## Abstract

The visual system is characterized by a strict topographic organization from the retina towards multiple layers of synaptic integration. Recent studies in Drosophila have shown that in the transition from the optic lobes to the central brain, due to convergence of columnar neurons onto optic glomeruli, distinct synaptic units employed in the computation of different visual features, the retinotopic representation is lost in these circuits. However, functional imaging revealed a *spatial representation of visual cues* in the Drosophila central complex, raising the question about the underlying circuitry, which bypasses optic glomerulus convergence.

While characterizing afferent arborizations within Drosophila visual glomeruli, we discovered a spatial segregation of topographic and non-topographic projections from distinct molecular classes of medulla projection neurons, *medullo-tubercular* (*MeTu*) neurons, into a specific central brain glomerulus, the *anterior optic tubercle* (*AOTu*). Single cell analysis revealed that topographic information is organized by ensembles of MeTu neurons (type 1), forming parallel channels within the AOTu, while a separate class of MeTu neurons (type 2) displays convergent projection, associated with a loss of spatial resolution. MeTu afferents in the AOTu synapse onto a matching topographic field of output projection neurons, these *tubercular-bulbar* (*TuBu*) neurons relay visual information towards dendritic fields of central complex ring neurons in the bulb neuropil. Within the bulb, neuronal proximity of the topographic AOTu map as well as channel identity is maintained despite the absence of a stereotyped map organization, providing the structural basis for spatial representation of visual information in the central complex (CX). TuBu neurons project onto dendritic fields of efferent ring neurons, where distinct sectors of the bulb correspond to a distinct ring domain in the ellipsoid body. We found a stereotypic circuitry for each analyzed TuBu class, thus the individual channels of peripheral topography are maintained in the central complex structure. Together with previous data showing rough topography within the lobula AOTu domain, our results on the organization of medulla projection neurons define the AOTu neuropil as the main relay station for spatial information from the optic lobes into the central brain.

## Introduction

Most flying diurnal insects rely on optic cues for accurate maneuvering, which requires appropriate processing of various visual stimuli (Egelhaaf & Kern 2002). Quick decisions whether to veer away or approach an object as well as being able to orientate within a three-dimensional environment are key tasks for their survival.

Research on neural networks at the periphery and in the central brain of a large variety of insect species, and foremost in Drosophila, has provided considerable insights into information processing beyond the compound eyes (Borst 2014; Silies et al. 2014; Behnia & Desplan 2015). While the resolution of an insect compound eye does not match that of a vertebrate retina (Kirschfeld 1976), neural substrates for the internal representation of the surrounding spatial world have been well documented (ref). Functional studies especially in flies have attributed distinct structures of the optic system dealing with different tasks of visual perception (Fisher et al. 2015; Schnell et al. 2010; Bahl et al. 2015). Of special interest has been the central complex (CX), a structure of interconnecting neuropils located at the midline of the protocerebrum. Its various functions comprise higher locomotor control, integration of sensory input and memory formation (Strauss 2002).

The CX plays a considerable role in visual information processing in insects, and neural pathways connecting the central body (CB), the largest part of the CX, with the optic lobes have been studied in various insect classes. Recent studies, using the variety of genetic tools in Drosophila, describing visual pattern memory (Liu et al. 2006), flight-dependent visual responses (Weir & Dickinson 2015) and visual landmark recognition (Seelig & Jayaraman 2015) suggest a substantial role of the central body for orientation in space and object recognition. While the neuroarchitecture of the CX contains aspects of a topographic map (Lin et al. 2013), the circuits and structural features that relay spatial information from the optic lobe into the CX are unknown.

Visual information is conveyed into the EB via distinct classes of R neurons, which form the ring-like layers of the EB (Hanesch et al. 1989; Wolff et al. 2015). Afferent neurons connect with R neurons, forming distinct synaptic units (microglomeruli) in the bulb, a neuropil adjacent to the EB, and formerly referred to as the lateral triangle (Ito et al. 2014a). These connections are distributed retinotopically, as they correlate to small receptive fields on the ipsilateral side (Seelig & Jayaraman 2013; Omoto et al. 2017). How spatial information is transmitted by optic lobe-descending neurons has so far not been investigated, however, a circuitry, involving two synaptic neuropils connecting the medulla with the EB, has been described from various insect taxa. A recent analysis of this pathway for Drosophila (Omoto et al. 2017) revealed distinct classes of medulla descending neurons (MeTu neurons) innervating the anterior optic tubercle (AOTu), a ventrolateral neuropil and a component of the set of ‘optic glomeruli’ in the central brain of insects. Optic glomeruli are characterized as synapse dense units of the ventrolateral neuropils, namely the AOTu, the ventrolateral protocerebrum (VLP) and the posteriorlateral protocerebrum (PLP) (Ito et al. 2014b; Otsuna & Ito 2006a; Panser et al. 2016). The majority of optic glomeruli are exclusively innervated by lobula columnar (LC) neurons (Otsuna & Ito 2006a). While recent functional studies in Drosophila revealed that distinct visual features are encoded in various optic glomeruli, e.g. small object detection, escape induction and reaching (Keleş & Frye 2017; Wu et al. 2016), in the majority of optic glomeruli spatial information is lost due to convergence and intermingling of LC neurons (Panser et al. 2016). In contrast, recent studies have shown that LC afferents in the AOTu display some rough spatial restriction along the dorso-ventral axis, providing the first indication of a topographic pathway into the central brain (Wu, Nern, W Ryan Williamson, et al. 2016).

Here, we show that medulla projection neurons in Drosophila form a stereotyped retinotopic map within the AOTu, which is spatially separated from LC representation. Interestingly, single receptive fields in the medulla diverge in multiple parallel visual channels, that are maintained in the subsequent synaptic bulb neuropil. Within the bulb, topographic channels connect with distinct receptive fields of central complex ring neurons.

Based on these data we are proposing a model in which the AOTu defines a central relay station for topographic information, organized in multiple parallel channels for visual object recognition.

## Results

### Distinct types of afferent arborization within optic glomeruli

Optic and olfactory glomeruli are prominent neuropil structures in different regions of the adult Drosophila brain, with olfactory glomeruli concentrated within the antennal lobes (AL) of the deutocerebrum, while the three neuropils containing optic glomeruli, the PLP, PLVP and AOTu (Fig. 1A), are part of the protocerebrum. To determine the structural similarities in connectivity of olfactory and optic glomeruli, we first analysed the arborization pattern of their afferent fibers. Olfactory glomeruli are characterized by a sensory class-specific convergence of afferent axons, thereby representing unique odorant receptor identities in the central brain (Laissue & Vosshall 2008) (Fig. 1B). Within each olfactory glomerulus, single sensory axon terminals arborize throughout the neuropil volume and axon terminal branches of all converging neurons broadly overlap and tightly intermingle (Hummel et al. 2003) (Fig. 1C).

**Fig.1:**
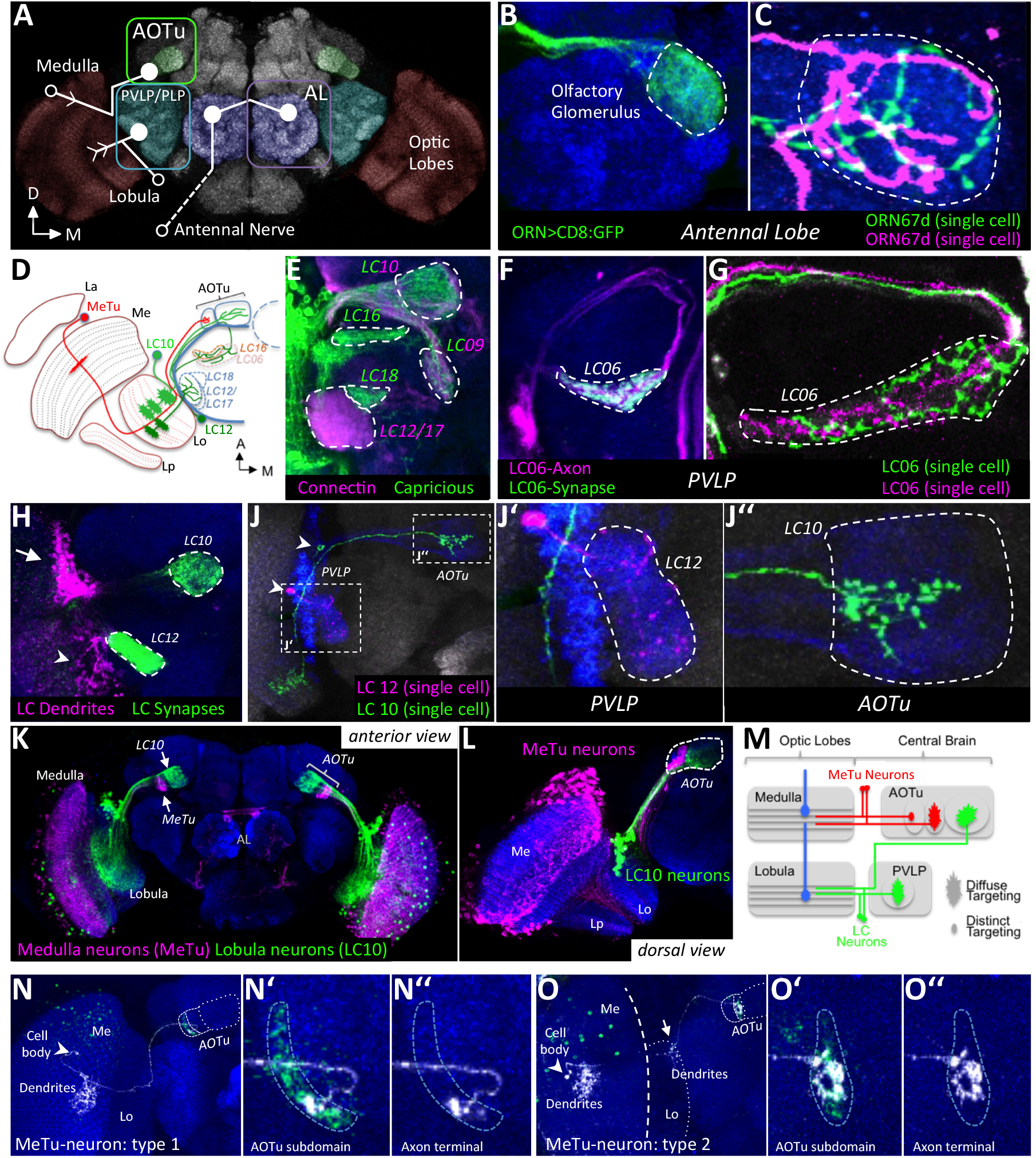
Organization of afferent projections within olfactory and optic glomeruli. If not otherwise mentioned, the frontal plane is shown. Names of glomeruli are written in italics. **A)** Overview of neuropils in the brain containing sensory glomeruli. Three pathways are shown, connecting medulla, lobula and antenna with their respective target neuropils (for clarity, lobula-AOTu connections are not drawn). Open circles represent the position of the cell body, closed circles a target glomerulus and arrows indicate dendritic arborizations. The antennal appendage is not part of the depicted brain. AL, antenna lobes; OL, optic lobes; AOTu, anterior optic tubercle; PVLP, posterior ventrolateral protocerebrum; PLP, posterior lateral protocerebrum; **B, C)** In the AL, axon terminals of OR67d neurons are branching throughout their target glomerulus DA1 and intermingle with each other. **D)** Schematic figure of visual projection neurons (VPNs) contributing to optic glomeruli (horizontal section). A subset of the >20 optic glomeruli (the AOTu and five representatives in the PVLP) are shown. Innervation of lobula layers is exemplified by one LC10 and one LC12 neuron. La, lamina; Lo, lobula; Lp, lobula plate. **E)** Optic glomeruli express different combinations of cell-adhesion molecules. **F)** LC06 neurons contribute to a characteristic optic glomerulus in the PVLP. The pre-synaptic region is visualized by GFP. **G)** Two LC06 neurons innervate the complete glomerulus and are not restricted to certain regions within the synaptic unit. **H)** LC10 and LC12 neurons are labeled by the same reporter line. Somato-dendritic and pre-synaptic compartments are visualized by specific markers. Cell bodies of LC10 are located at the dorsal (arrow), those of LC12 on the ventral side (arrowhead) of the lobula neck. **J-J’’)** Single cell morphologies of LC10 and LC12 neurons. While LC12 neurons branch throughout their target glomerulus (J’), LC10 neuron terminals are dorso-ventrally restricted within the LU (J’’). Position of cell bodies are indicated by arrowheads. **K, L)** The AOTu consists of compartments that are innervated by either medulla neurons (MeTu) or lobula neurons (LC10). Additionally, some cells in the medulla express GFP under LexA control. Not all MeTu neurons are labeled with this expression line. **M)** Schematic pathways and sizes of axonal terminals of afferent neurons innervating the AOTu and PVLP. Afferent medulla innervation indicated by blue neurons. MC61 innervation of lobula layers (compare (O)) and the PLP neuropil are not shown. **N-O’’)** Single cell labeling reveals the projection route and axonal terminal size of MeTu neurons. Different subtypes of MeTu neurons can be determined by the position of their respective target region, size of terminal arborization and whether the lobula is innervated in addition (arrowhead in (O)). Arrowheads indicate position of cell bodies. Genotypes: B) *OR47d::GFP* C) *hs-mFlp5; OR67d>FLYBOW1.1* E) *Caps>GFP (MARCM)* F) *R41C07>mCherry, >syt::GFP* G) *hs-mFlp5; R41C07>FLYBOW1.1* H) *R35D04>DenMark, >syt::GFP* J) *hs-mFlp5; R35D04>FLYBOW1.1 K,L) *VT29314-LexA>GFP; R44A03>mCherry** N) *hs-mFlp5; R52H03>FLYBOW1.1* O) *hs-mFlp5; R20B05>FLYBOW1.1*

Optic glomeruli in Drosophila receive their main neuronal input from lobula columnar (LC) neurons connecting the optic lobes with the PLP/PLVP region in the ventrolateral brain region (Otsuna & Ito 2006b) (Fig. 1D). In addition, the AOTu, located dorsally in respect to the PLP/PLVP region, is a major optic glomerular neuropil with afferent input via the anterior optic tract (Fischbach & Lyly-Hünerberg 1983; Otsuna & Ito 2006b; Panser et al. 2016; Omoto et al. 2017)(Fig. 1J). Using the FlyLight and Vienna Tiles collection of specific neural expression lines (Jenett et al. 2012; Kvon et al. 2014), a variety of LC neuron types could be identified and their class-specific segregation into single optic glomeruli visualized (Costa et al. 2016; Panser et al. 2016) (Fig. 1F-L). In addition to their morphological pattern and spatial position within the PLP/PLVP region, optic glomeruli, similar to olfactory glomeruli in the antennal lobe (Hong et al. 2012; Hong et al. 2009), can be classified according to the expression of different types of cell surface molecules (Fig. 1E for the LRR-type proteins Connectin and Capricious and data not shown).

To determine the afferent arborization within optic glomeruli, single cell clones for various LC neuron types were generated (Fig. 1G, H). Similar to olfactory sensory neurons (OSN) axon terminals, each LC axon ramifies throughout a single optic glomerulus and all neurons of the same LC class converge onto a common glomerular space (Fig. 1G, H), confirming recently published data (Wu et al., 2016). In contrast to the homogeneous arborization pattern within synaptic glomeruli of the PVP/PLVP neuropil, a more diverse pattern of afferent innervation was observed for the AOTu area, which can be subdivided into a medially located large unit (LU), which has been named AOTUm in (Omoto et al. 2017), and a lateral, small unit (SU) (Fig.1D, K). Systematic characterization of a large collection of AOTu-specific expression lines showed that the LU and SU regions represent the target fields of two distinct optic neuropils, in which the LU receives input from the lobula (LC neurons), whereas the SU is targeted exclusively by medulla projection neurons (MeTu neurons, Panser et al. 2016, Moto et al. 2017) (Fig. K-M, and see below). Afferent innervation of the LU arborizes through large parts of the glomerular neuropil, although some spatial enrichment in the dorsal versus ventral glomerulus can be detected (Wu, Nern, W. Ryan Williamson, et al. 2016) (Fig. 1J’’). In contrast, afferent patterns in the SU ranges from broad arborization in a region adjacent to the LU (MeTu-type 2 neurons) to a highly restricted termination in more lateral regions (MeTu-type 1 neuron, Fig. 1N, O), which suggests different levels of spatial representation within the AOTu, analyzed in more detail below.

In summary, regarding neuron-type specificity and the spatial organization of afferent fiber projections within the synaptic target field, the majority of optic glomeruli recapitulate the principle of axonal convergence in which terminal arborizations of all neurons of a given type tightly intermingle with no signs of topography, indicating a loss of spatial information. In contrast, the AOTu neuropil contains a range of afferent targeting patterns with various degrees of intraglomerular spatial restriction and nonoverlapping afferent terminals, which makes this pathway an interesting candidate for topographic representation of visual information in the central brain.

### Sub-domain organization of the AOTu

To determine how the organization of the AOTu neuropil correlates with the pattern of afferent innervation, we first visualized glial membranes combined with the neuropil epitope N-Cadherin (Fig. 2A, B). In addition to the distinction between the LU and SU regions of the AOTu, a further subdivision of the SU region with multiple domains along the medial-lateral axis could be detected, whereas the LU neuropil possessed a homogeneous appearance (Fig. 2A, B).

**Fig.2:**
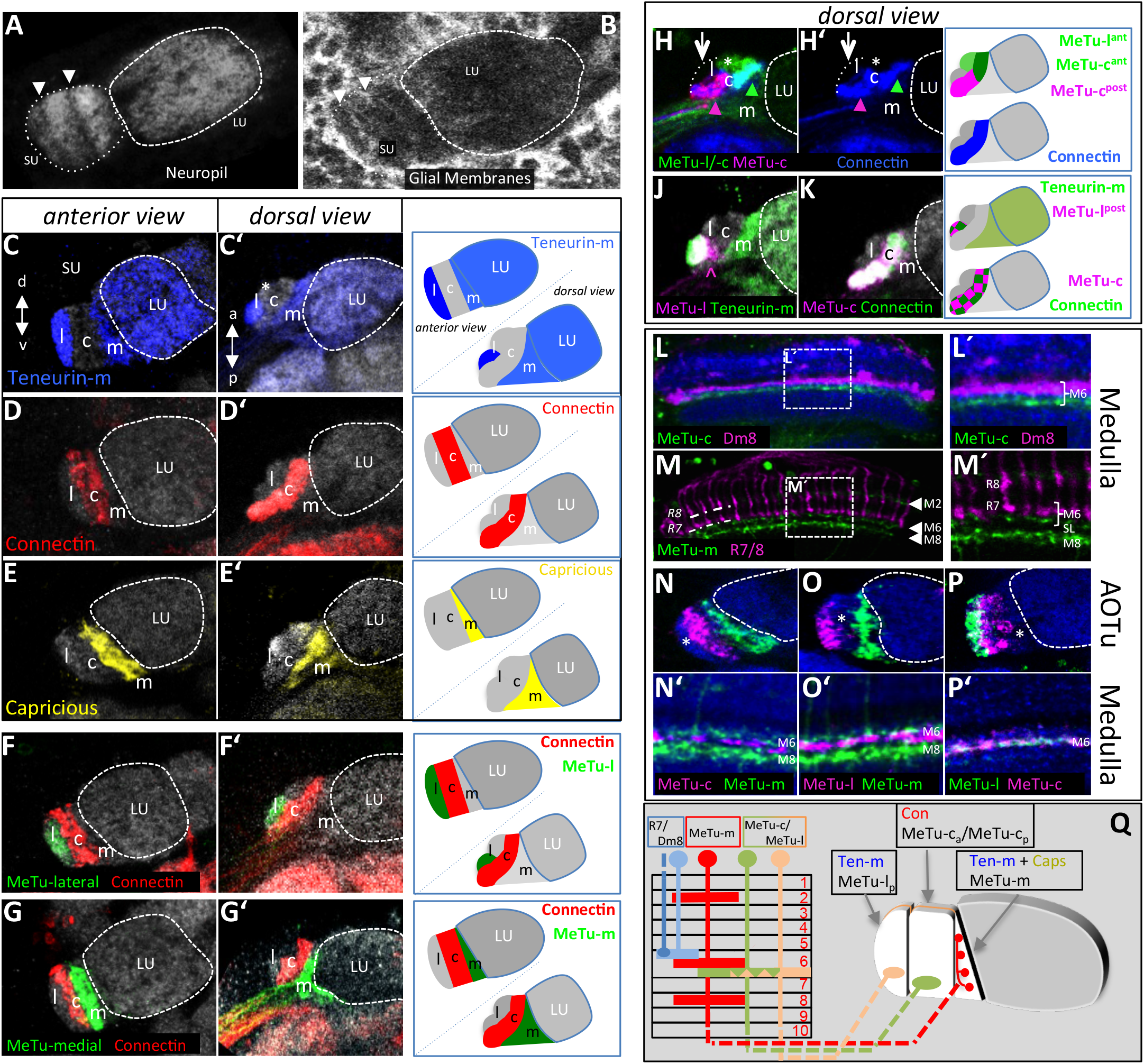
Classification of MeTu-neuron subtypes. The AOTu can be divided into a large unit (LU), innervated by afferent LC10 neurons, and a small unit (SU), innervated by afferent MeTu neurons. The SU is further subdivided and different types of MeTu neurons can be distinguished, innervating separated regions within the SU. **A)** The division of the SU can readily be observed with neuropil markers (anti-CadN). Arrowheads indicate borders of domains or subdomains. In contrast, the adult LU has a uniform appearance. **B)** Glial labeling reflects the compartmentalization of the AOTu (arrowheads), while no apparent compartments are visible in the LU. **C-E’)** Each SU-domain can be uniquely characterized by a combination of three cell-adhesion molecules (CAMs). **C, C’)** Ten-m is strongly expressed in the lateral domain, with lower intensity in the medial domain and the LU. The dorsal view (C’), shows that the lateral domain is further divided into an anterior, Ten-m negative (asterisk) and a posterior, Ten-m-positive compartment. **D, D’)** Con expression is specific for and defines the central domain. **E, E’)** GFP expressed under the control of Caps-Gal4 marks the medial domain. **F-G’)** Domain borders defined by CAM-expression are respected by MeTu-neurons. Gal4-labeled neurons innervate the lateral (F-F’) or medial domain (G-G’), without overlapping the central, Con-positive domain. **H-K)** A combination of immunochemistry and transgenic expression reveals a further division of the lateral and central domain into anterior and posterior compartments. **H-H’)** Combining LexA-(green) and a Gal4-(magenta) expression lines reveals a subdivision of the central domain. Anti-Con labeling is used to visualize the complete central domain. A subset of neurons labeled by the LexA-line also innervates the anterior lateral domain (asterisk). **J)** Labeled neurons innervate exclusively the posterior part of the lateral domain, which is also defined by Ten-m expression. The asterisk marks the anterior, unlabeled part of the lateral domain. Note that neurons innervating the lateral domain form a turn upon arriving at the AOTu, thus the arrowhead marks turning axons and is not part of the target field. **K)** The complete central, Con-positive, domain is labeled by this expression line. **L-M’)** The outer medulla is innervated by both types of MeTu neurons. Only MeTu-type 2 neurons form in addition dendritic branches in the inner medulla. **L, L’)** Dendrites of MeTu-c neurons are exclusively innervating the proximal M6 layer. There is no direct contact between the color-vision circuit and MeTu-neurons. **M, M’)** Three medulla layers are innervated by MeTu-m neurons (arrowheads). Photoreceptor neurons R7 and R8 are labeled with anti-chaoptin (24B10). SL, serpentine layer. **N-P’)** Separation and overlap of MeTu neurons in the medulla and SU. MeTu-c and MeTu-l neurons do not share an M6 sub-layer with MeTu-m neurons. Asterisks indicate the respective unlabeled domain. In the upper row (N, O, P) the AOTu and in the lower row (N’, O’, P’) the medulla of the same specimen are shown, respectively. **N-N’)** The central and medial domain are labeled. **O-O’)** Neurons innervating the lateral and medial domain are labeled. **P-P’)** MeTu-c and MeTu-l neurons are separated in the SU, but share the same layer in the medulla. **Q)** Overview of innervation patterns of MeTu neurons in medulla and SU. Within the M6-layer, three separate sub-layers can be distinguished, the proximal two innervated by MeTu neurons, with no direct connection to photoreceptor neurons R7, which terminate in the distal part of M6. Genotypes: B) *Repo>GFP* E) *Caps>GFP* F, F’) *R85F05>GFP* G, G’) *R20B05-LexA>GFP* H) *R25H10>mCherry; R67C09-LexA>GFP* J) *R85F05>GFP* K) *R44A03>GFP* L) *R56F07>mCherry, Dm8-LexA>GFP* M) *R20B05-LexA>GFP*. N,N’) *R56F07>mCherry, R20B05-LexA>GFP* O,O’) *R56F07>mCherry, R20B05-LexA>GFP* P,P’) *R44A03>mCherry, R94G05-LexA>GFP*

We next analyzed how the SU neuropil domains match with the afferent innervation from the medulla (Fig. 2C-K). Based on the axon arborization pattern from 13 independent expression lines we could distinguish three non-overlapping populations of MeTu neurons with domain-specific SU innervation, which were named accordingly MeTu-lateral (-l), MeTu-central (-c) and MeTu-medial (-m) (compare Fig. 2N-O). This morphological neuropil organization was further supported by the combinatorial expression pattern of various neuronal cell adhesion molecules. For example, the synaptic cell adhesion molecule Teneurin-m is expressed broadly in the AOTu neuropil with the exception of the central SU (SU-c) and the anterior SU-l domains (Fig. 2C). On the other hand, the LRR-type adhesion molecules Connectin and Capricious are specifically expressed in the SU-c and SU-m domains, respectively (Fig. 2D, E, F, G). Furthermore, detailed analysis of molecular markers in combination with MeTu expression lines revealed an additional subdivision of the SU-l domain along the anterior-posterior axis (SU-l_a_ vs SU-l_p_, Fig. 2C’, F’), which was not detected in the other SU domains or the LU neuropil (Fig 2C’, D’, E’ G’). Furthermore, by combining two expression lines (Gal4 + LexA system) we additionally discovered a division of the SU-c (Fig. 2H). To show that this subdivision is in fact a property of MeTu neurons, an expression line was co-labeled with a molecular marker (Fig. 2J, H’). In contrast there are expression lines, which label a complete domain (Fig. 2K). In summary, the AOTu neuropil is organized in previously undescribed morphological and molecularly defined subunits of afferent innervation.

To get further insights into the different MeTu neuron populations, we analyzed their layer-specific dendritic arborization in the medulla neuropil, which is organized into distinct synaptic layers (Fig. 2L-P’). Interestingly, all three MeTu classes innervate layer 6 of the medulla neuropil, the main synaptic region for R7 photoreceptor input. For MeTu-l and MeTu–c neurons, layer 6 is the only site of dendritic field positioning (Fig. 2L, P’), In contrast, MeTu-m neurons contain dendritic arborizations in two additional medulla layers both proximal and distal to layer 6, most likely layer 2 and layer 8 ((Fig. 2M, M’, N’, O’). Further characterization of MeTu dendrites within layer 6 revealed a non-overlapping arrangement with the R7/Dm8 synaptic area (Fig. 2 L, M). In addition, within layer 6 MeTu-m dendrites clearly segregate from the MeTu-l/-c dendritic location (Fig. 2N’,O’), which allows to define three distinct sub-layers (R7/Dm8, MeTu-m, MeTu-l/-c) within medulla layer 6 (Fig. 2Q). Thus, the AOTu receives direct input from the medulla via various types of MeTu neurons, which differ in their dendrite position, axon targeting and molecular identity (summarized in Fig. 2Q).

### Topographic organization of AOTu afferents

We next determined how AOTu afferent patterning correlates with the MeTu-neuron ensembles of distinct molecular identity. Single cell analysis revealed a stereotyped pattern of MeTu-neuron innervation within the different SU domains, in which each MeTu-neuron sends an axon to only one of the three SU subdomains (Fig. 3A-C). For MeTu-l and MeTu-c neurons, a spatially restricted afferent termination can be observed in the respective SU compartments (Fig. 3A, B). In contrast, MeTu-m afferent arborization extends through a large part of the compartment, which resembles the projection of LC10 neurons in the adjacent LU (Fig. 1J). These results show that the morphological and molecular subdivisions of the AOTu correlate with a distinct connectivity pattern of MeTu-neurons. MeTu-m neurons with dendritic arborizations in multiple medulla layers show axonal convergence similar to the majority of optic glomeruli innervated by LC neurons. In contrast, MeTu-l and MeTu-c neurons, the two medulla projection neuron classes with restricted axon terminals in the AOTu, display identical dendrite layer positioning in layer 6 but show axon segregation in distinct SU domains, indicating parallel visual channels into the central brain.

**Fig.3:**
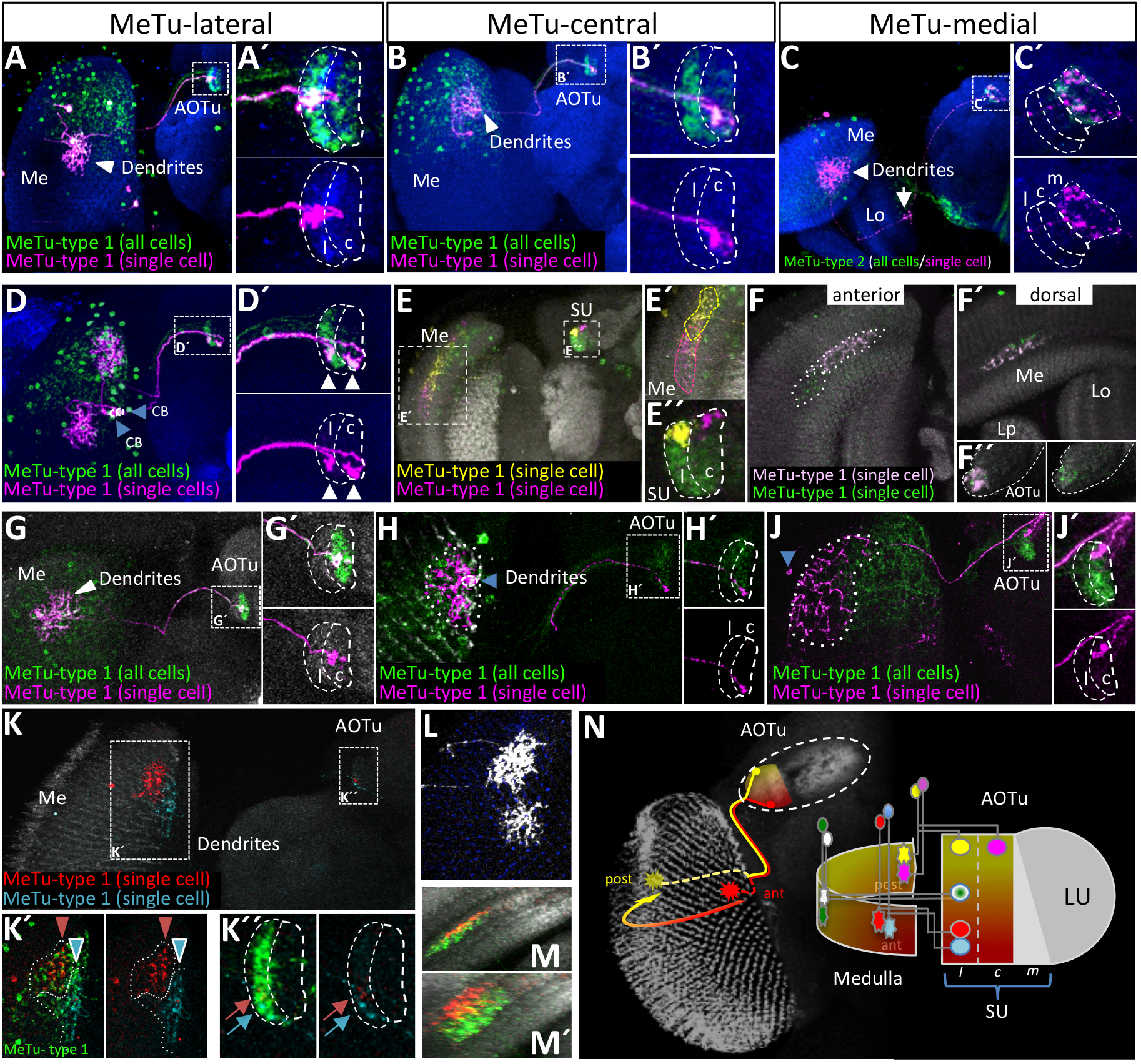
Topographic organization of AOTu projections. **A-C’)** FLYBOW-labeling of MeTu-neurons innervating the respective three domains of the SU. Arrow in (C) indicates innervation of the lobula by MeTu-m neurons. Note that the optic lobe is tilted dorsally. **D, D’)** Two neighboring cells (blue arrowheads) innervate different positions within the dorsal medulla and target the lateral and the central SU-domain, respectively (white arrowheads). CB, cell body. **E, E’)** MeTu-type 1 neurons at the posterior edge of the medulla exclusively target dorsal areas of the AOTu. Dendritic fields overlap among MeTu neurons. The lateral domain (yellow neuron) and the central domain (magenta neuron) are innervated. **F-F’’)** Dorso-ventral positions in the medulla do not correlate to topographic projections in the AOTu. MeTu-c neurons with the same A-P position in the medulla target into an overlapping region in the central SU-domain. Different angles of the same brain are shown. **G-J’)** Topographic projections of MeTu-c neurons: Central dendritic fields in the medulla are maintained in the AOTu (G-G’), anterior dendritic positions in the medulla translate to a ventral targeting (H-H’), while posterior medullar innervation correlates to a dorsal position (J-J’). The dimension of dendritic fields does not correlate to the size of the innervated target area: In (H), 6 columns in A-P axis and 5 columns in D-V axis are innervated. In (J), the dendritic field of one cell stretches about 1/3rd in A-P axis and almost over the complete dorsal half of the medulla. Blue arrowheads indicate the cell bodies. **K-K’’)** Dendritic fields at the anterior border of the medulla correlate with a ventral innervation in the AOTu. The red neuron – in comparison to the cyan one - being more posterior in the medulla translates to a more dorsal position in the AOTu. **L)** The size of dendritic fields varies amongst MeTu-l neurons. **M, M’)** Overlap of dendritic fields in two MeTu-l neurons. Different angles of the same brain are shown. **N)** Overview of the FLYBOW-pairs shown in E, F, and K (colors accordingly) and a model of topographic correlations between medullar dendritic fields and the SU axis of innervation.Genotypes: A), B) *hs-mFlp5; R52H03>FLYBOW1.1* C) *hs-mFlp5; R20B05>FLYBOW1.1* D) *hs-mFlp5; R52H03>FLYBOW1.1* E) *hs-mFlp5; R52H03>FLYBOW1.1* F) *hs-mFlp5; R52H03>FLYBOW1.1* G-J) *hs-mFlp5; R56F07>FLYBOW1.1* K) *hs-mFlp5; R85F05>FLYBOW1.1* L) *hs-mFlp5; R85F05>FLYBOW1.1* M) *hs-mFlp5; R85F05>FLYBOW1.1*

All types of MeTu-neurons have dendritic fields that are restricted to the dorsal half of the medulla neuropil (Fig 3A-E). Although MeTu-neurons cover multiple medulla columns, dendritic field size of individual MeTu-neurons - within and between MeTu classes - can vary substantially (Fig. 3L). Random single cell clones revealed two main features in the spatial organization of MeTu projections: 1) MeTu-neurons of the same type (MeTu-l/-l or MeTu-c/-c) with neighboring dendritic fields project to adjacent positions in the corresponding SU domain (Fig. 3K); 2) MeTu-neurons of different types (MeTu-l/-c) with overlapping dendritic fields in the medulla project to the same position along the d-v axis in adjacent SU domains (Fig. 3E).

To determine if the spatial patterning of MeTu-l and - c projections within their corresponding SU domain reflects a retinotopic representation of the medulla columnar field, we compared the relative position of dendrites and axon terminals in the medulla and AOTu respectively (Fig. 3G-J). For both MeTu-l and MeTu-c neurons we could observe a strict correlation of the dendrite/axon position between the anterior-posterior (a-p) axis in the medulla and the dorso-ventral (d-v) axis in the AOTu (Fig. 3N, n=35). Here, all MeTu-l/-c neurons with dendrites at the anterior edge of the medulla neuropil target to the most ventral position in the corresponding SU domain whereas neurons with dendrites at the posterior medulla edge connect to a dorsal SU position (Fig. 3H, J). Furthermore, all MeTu cell clones with dendrites in the medial medulla region target to a medial position in the AOTu. In contrast, for the spatial arrangement of dendrites along the d-v axis of the medulla we did not detect a distinct MeTu-l/-c axon targeting pattern in the SU domains, in which the neurons with adjacent dendrites that span the medulla field along the d-v axis converge onto the same SU target region (Fig. 3F). Furthermore, single MeTu-l and MeTu-c neurons with overlapping dendritic fields target their axons to comparable d-v positions in adjacent SU domains (Fig. 3E). In contrast to the defined connectivity of MeTu-l/-c, no obvious spatial correlation between dendrite position and SU-m targeting could be detected for MeTu-m neurons. These data support a topographic representation of visual information from the dorsal retina along the anterior-posterior axis, providing the structural basis for the transmission of spatial information in the central brain of Drosophila.

### AOTu projection neurons maintain domain identity and visual topography

If the AOTu defines a relay station of spatial information from the optic lobes to the central brain, we would expect a matching patterning between AOTu output neuron and MeTu projections along the d-v axis of the SU domains.

We identified a large set of expression lines for AOTu projection neurons targeting the bulb region (TuBu neurons, Omoto et al. 2017), which show a domain-specific restriction of their dendritic fields, corresponding to the SU-l, - c and - m domains, therefore classified as TuBu-l, - c & - m neurons (Fig. 4B, G; compare (Omoto et al. 2017)). In single cell clones, the size of TuBu dendrite fields matches the corresponding MeTu axon terminal arborizations, in which TuBu neurons of the SU-l and - c domains show highly restricted dendritic arbors whereas TuBu-m neurons can also form broad dendritic fields (Fig. 4H, J). In addition, the average number of TuBu neurons covering a SU domain along the d-v axis corresponds well to the average number of MeTu neurons representing the ant-post medulla axis (between 8-12 neurons for different classes), indicating a point-to-point representation rather than convergence. To test if the spatial overlap of MeTu axon terminals and TuBu dendrites indicate synaptic connections we employed the syb-GRASP technique, which visualizes synaptic contacts based on split-GFP reconstitution following vesicle release (Karuppudurai et al. 2014). Here, GRASP analysis between presynaptic MeTu neuron subtypes and various sets of TuBu neurons revealed a strict matching of synaptic partners within, but not across SU domains (Fig. 4C-D), supporting the model that the SU molecular domain identity corresponds to synaptic recognition.

**Fig.4:**
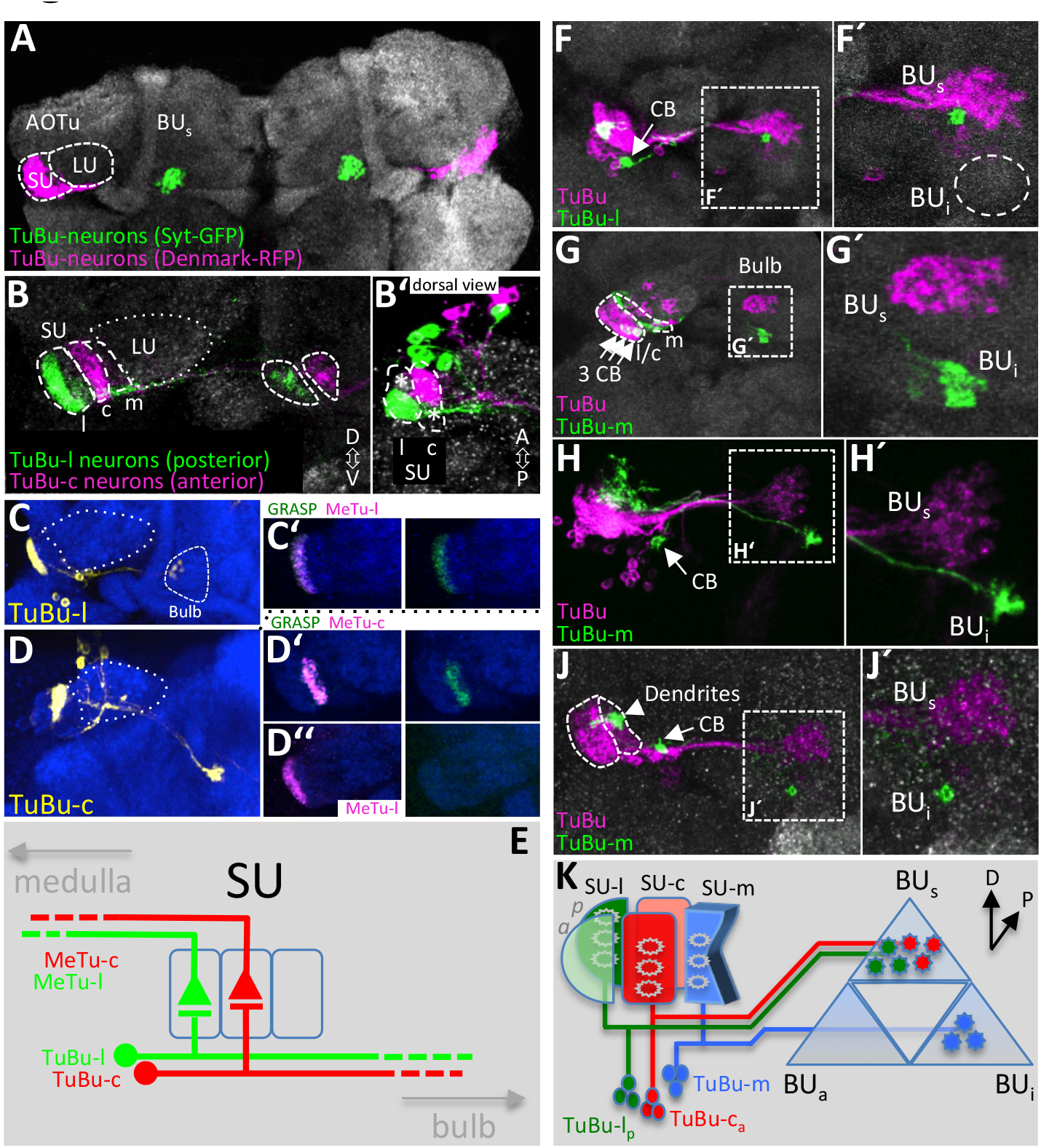
Bulb-innervating neurons descending from the AOTu maintain domain identity. View is anterior in all images, if not otherwise mentioned **A)** The bulb receives input from all three SU-domains. **B, B’)** Within the bulb, terminals of TuBu-l and TuBu-c neurons are spatially separated. Asterisks mark the anterior lateral and the posterior central SU-domain, which are not labeled by these expression lines. **C-D’’)** Pre postsynaptic matching of domain-specific expression lines in the SU revealed by synGRASP. The anti-GFP antibody exclusively detects the presynaptic (GFP1-10) construct, expressed under Gal4-control. **C-C’)** Positive GRASP-signal in combination of MeTu-l and TuBu-l neurons. **D-D’’)** TuBu-c neurons are synaptic partners of MeTu-c neurons (D’). Across neighboring domains no direct synaptic connections are formed between MeTu and TuBu-neurons (D’’). **E)** Afferent medulla neurons and efferent TuBu-neurons form circuits in their respective SU-domains. **F-J’)** FLYBOW-labeling with a reporter line for the majority of TuBu neurons. Intensity of the default marker (magenta) is very low for TuBu neurons targeting the BUi. CB, cell body. **F, F’)** TuBu-l neurons cover restricted areas in both dendritic and axonal terminals. Dashed circle indicates the approximate position of the BUi. **G)** TuBu-m neurons innervate a ventral area in the bulb (BU_i_), separated from TuBu-l and TuBu-c neurons. Three TuBu-m neurons are GFP labeled. **H, J)** TuBu-m neurons are variable in arborization size in both AOTu and bulb, either restricted (J) or covering larger areas (H). **K)** Distribution of three TuBu-classes in the bulb. The remaining classes likely innervate separate areas of the BU_s_ Genotypes: A) *R86C02>DenMark, >syt::GFP,* B) *R64F06>mCherry, R25F06-LexA>GFP,* C) *R25F06-LexA>GFP*, C’) *R85F05>syb::spGFP1-10; R25F06-LexA>CD4::spGFP11*, D) *R64F06-LexA>GFP*, D’) *R56F07> syb::spGFP1-10; R64F06-LexA>CD4::spGFP11*, D’’) *R85F05>syb::spGFP1-10; R64F06-LexA>CD4::spGFP11*, F-J) *hs-mFlp5; R86C02>FLYBOW1.1*.

The axons of all TuBu neurons join a common fascicle which extends towards the central bulb neuropil region, where they segregate into distinct domains according to their AOTu domain identity (Fig. 4K; compare also Omoto et al. 2017): The TuBu-l and - c neurons terminate into adjacent regions in the superior bulb (BUs), whereas axons of TuBu-m neurons target into a distinct inferior bulb (BUi) area (Fig. 4F, G), demonstrating a spatially segregation of topographic and convergent visual pathways within the bulb. Using a collection of various TuBu expression lines we next analyzed the projection of single cell and small size clones regarding here spatial organization of dendritic and axons arborization.

To determine if the retinotopic representation in the AOTu is maintained in the TuBu connectivity within the bulb region, we compared the relative positions of dendrites in the SU with their axon termination in the bulb region of individual TuBu neurons. Furthermore, two-cell clones of MeTu-l neurons showed that adjacent dendritic positions in the AOTu are maintained within afferent domains of the bulb regions, although their relative position within the bulb area is variable (Fig. 5A, B). To further determine the spatial patterning of TuBu neurons we generated a series of single cell clones and compared the relative position of SU dendrites with axon termination in the bulb region of individual TuBu neurons (Fig. 5D). In contrast to the strict spatial correlation for MeTu neuron dendrite/axon position along the ant-post/d-v axis, for TuBu neurons the position of the dendritic field within the SU domain does not predict the site of axon termination within the respective bulb area (Fig. 5E, F). For example, independent TuBu-l single cell clones with dendritic fields in the dorsal SU domain show projections to the dorsal, ventro-lateral or ventral-medial bulb domain (Fig. 5F, left column). Similarly, the dorsal bulb region can receive TuBu innervation from neurons with dorsal, medial or ventral SU positions (Fig. 5F, right column). Together with the spatial proximity of axon targeting for adjacent TuBu neurons these data suggest that the topographic map of the AOTu translates into a circular representation with more dynamic positioning within the bulb.

**Fig.5:**
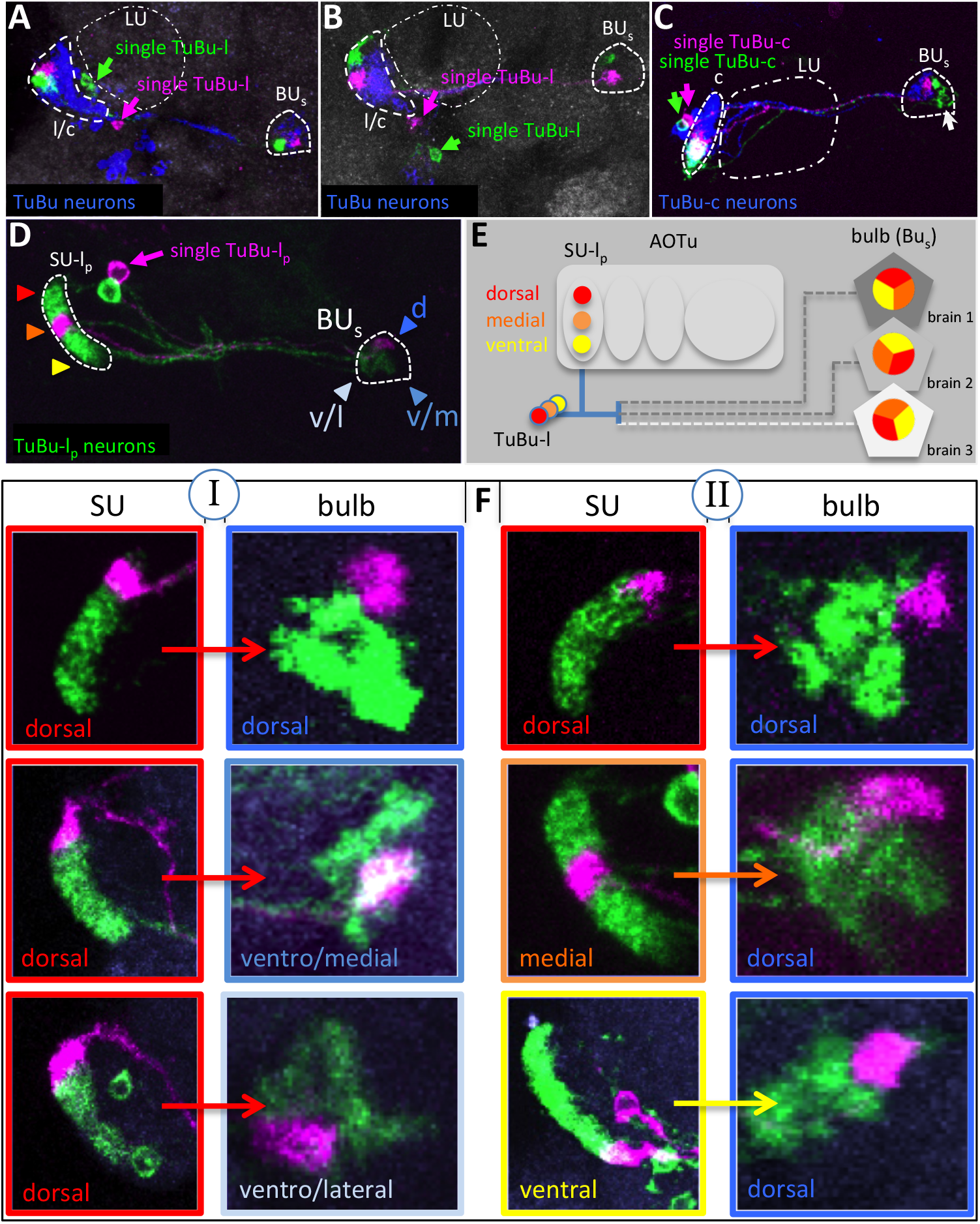
Variability of innervation patterns in descending TuBu neurons. **A-C)** Neighboring neurons maintain their proximity within the bulb, but there is variance in their orientation. In (A) and (B) TuBu neurons from all three domains are labeled, in (C) an expression line for the central domain is used. Approximate borders of the lateral/central SU domain, LU and superior bulb are indicated. **D)** FLYBOW-single cell clone (magenta) with an expression line specific for TuBu-l neurons. Arrowheads: color coding for dorso-ventral distribution in the AOTu and positions in the BUs (dorsal, ventro-lateral, ventro-medial), also used in the subsequent panels. **E)** TuBu-lp neurons innervate adjacent regions in the bulb, but there is no stereotypic orientation. **F)** No topographic correlation between dendritic position in the AOTu and target field in the bulb. Neurons with dorsal positions in the AOTu target to various positions within the lateral sector of the BUs (column I). Likewise, a similar position in the bulb gets innervation from all positions along the D-V axis in the AOTu (column II). Genotypes: A) *hs-mFlp5; R71E07>FLYBOW1.1*, B) *hs-mFlp5; R86C02>FLYBOW1.1*, C) *hs-mFlp5; R64F06>FLYBOW1.1*, D, F) *R25F06>FLYBOW1.1*

### Projections of AOTu domain identity onto ring neurons of the EB

Efferent neurons of the bulb region have been shown to target specific ring layers within the EB (R neurons). Co-labeling showed that dendrites of different R neuron types segregate into coherent, non-overlapping domains within the bulb neuropil (Fig. 6 G-J). To determine the matching of TuBu target domains and the spatial positioning of dendritic areas of R neuron types, we performed a series of co-labeling and GRASP studies (Fig. 6 A-F and data not shown), for which we received the most complete set of data for two TuBu-classes (TuBu-l_post_ & TuBu-c_ant_) in combination with R neurons of the BUs. The BUi is not targeted by TuBu-l or TuBu-c neurons (data not shown, compare (Omoto et al. 2017). We could identify a class-specific projection pattern, in which TuBu axons either fully target to a single R neuron type (Fig. 6A) or are only partially overlapping with dendritic fields of Ring neurons (Fig 6E). While both TuBu classes and three R neuron classes innervate the superior bulb, the observation that only for one combination a robust AOTu-CX circuit has been detected indicates a high specificity within a small part of the bulb. That no spatial overlap with R neurons could be detected for TuBu-c efferents, supports the view of a distinct pathway of information from the two topographic SU domains into the bulb (Fig. 6K).

**Fig.6:**
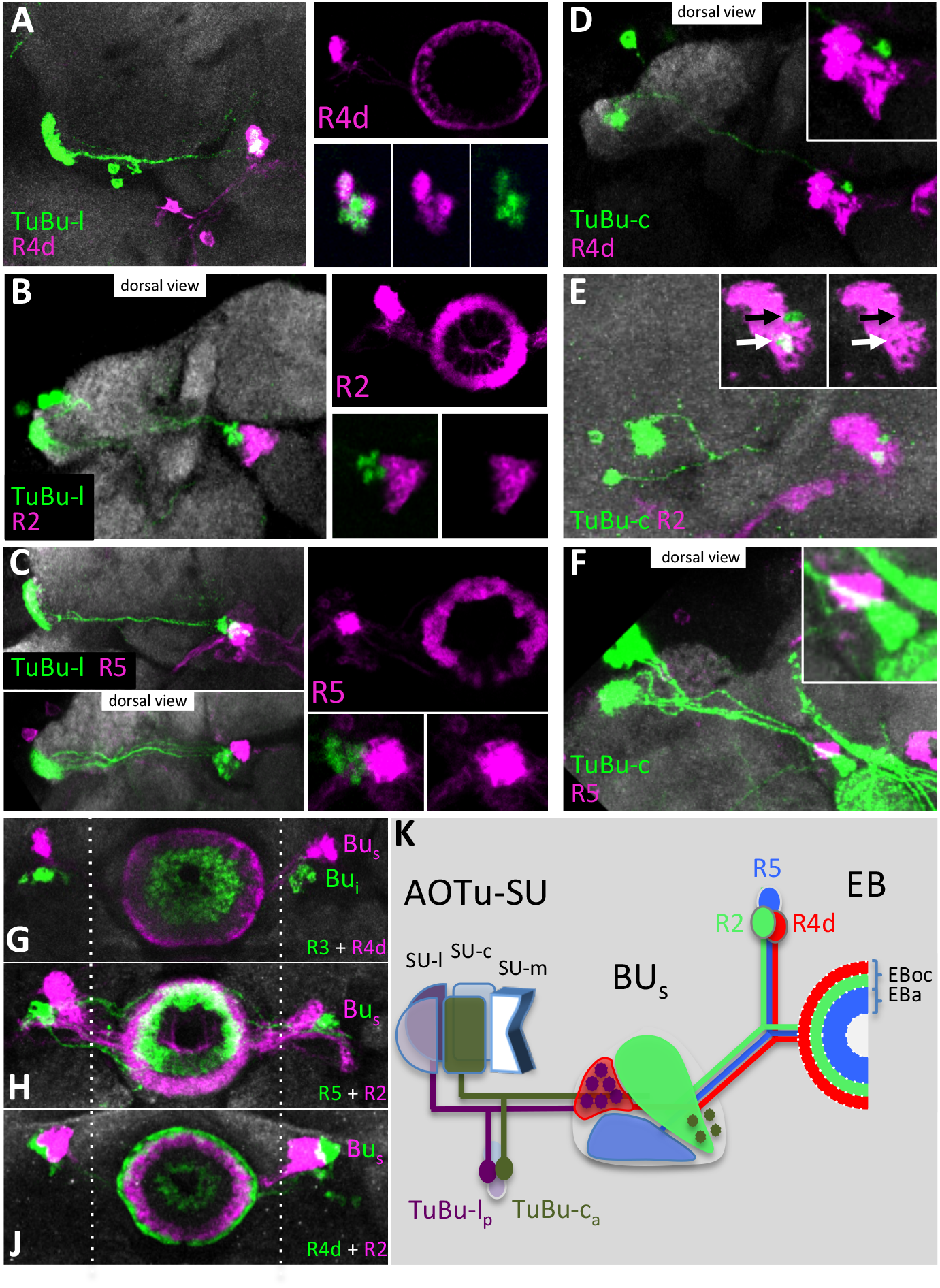
Distinct AOTu pathways connect with specific R neuron classes. **A)** In the BUs different TuBu classes connect to a set of R neurons. Two LexA expression lines label the posterior lateral domain and the anterior central domain of the SU, respectively. The BUa and BUi are not covered in this analysis. **B-C)** Co-labeling of R neurons reveals the coverage different fields within the BU. Genotypes: **E-H)** TuBu-lp neurons () exclusively contact with R4d neurons. Gal4 lines used for R neurons labeling: **J-L)** TuBu-c_a_ neurons (R64F06-LexA) partially overlap with R2 neurons. Gal4 lines same as in (E)-(G). **K)** Proposed segregation of visual information via the AVP (anterior visual pathway): Retinotopic arrangements in the medulla and AOTu are transferred to variable but distinct areas within sectors of the bulb.
Genotypes: B) *R12B01>mCherry; R14A12-LexA>GFP*, C) *EB1>mCherry; R48H04-LexA>GFP*, D) *EB1>mCherry; R85E07-LexA>GFP*, E) *R12B01>mCherry; R25F06-LexA>GFP*, (F) *EB1>mCherry; R25F06-LexA>GFP*, (G) *R49B02>mCherry; R25F06-LexA>GFP*, (H) *R14A12>mCherry; R25F06-LexA>GFP*, J-L) *R64F06-LexA>GFP*, Gal4-lines same as in (E-H).

## Discussion

Like various other sensory modalities for which spatial information is critical, neural circuits in the visual system are organized in a topographic fashion to maintain the activity pattern of peripheral photoreceptors along the visual pathways into the central brain. Although spatially-patterned visual stimuli induce neural activity in the central complex of the Drosophila brain (Seelig & Jayaraman 2013), the topographic pathway that translates peripheral information into central activity patterns remain elusive. Here we show that in the Drosophila central brain, optic glomeruli, relay stations of peripheral visual information, fall into two morphological types regarding their afferent arborization pattern: broad arborization of intermingling terminal axons or spatially-restricted, non-overlapping axon terminals. While the retinotopic representation within the lobula neuropil is lost in the broad arborization pattern of converging LC projection neurons onto the majority of optic glomeruli (Panser et al. 2016; Wu, Nern, W Ryan Williamson, et al. 2016; Keleş & Frye 2017), we could identify a unique spatial organization for the evolutionary-conserved output channel from the medulla, the medulla-AOTu fiber tract (Fig. 7).

**Fig.7:**
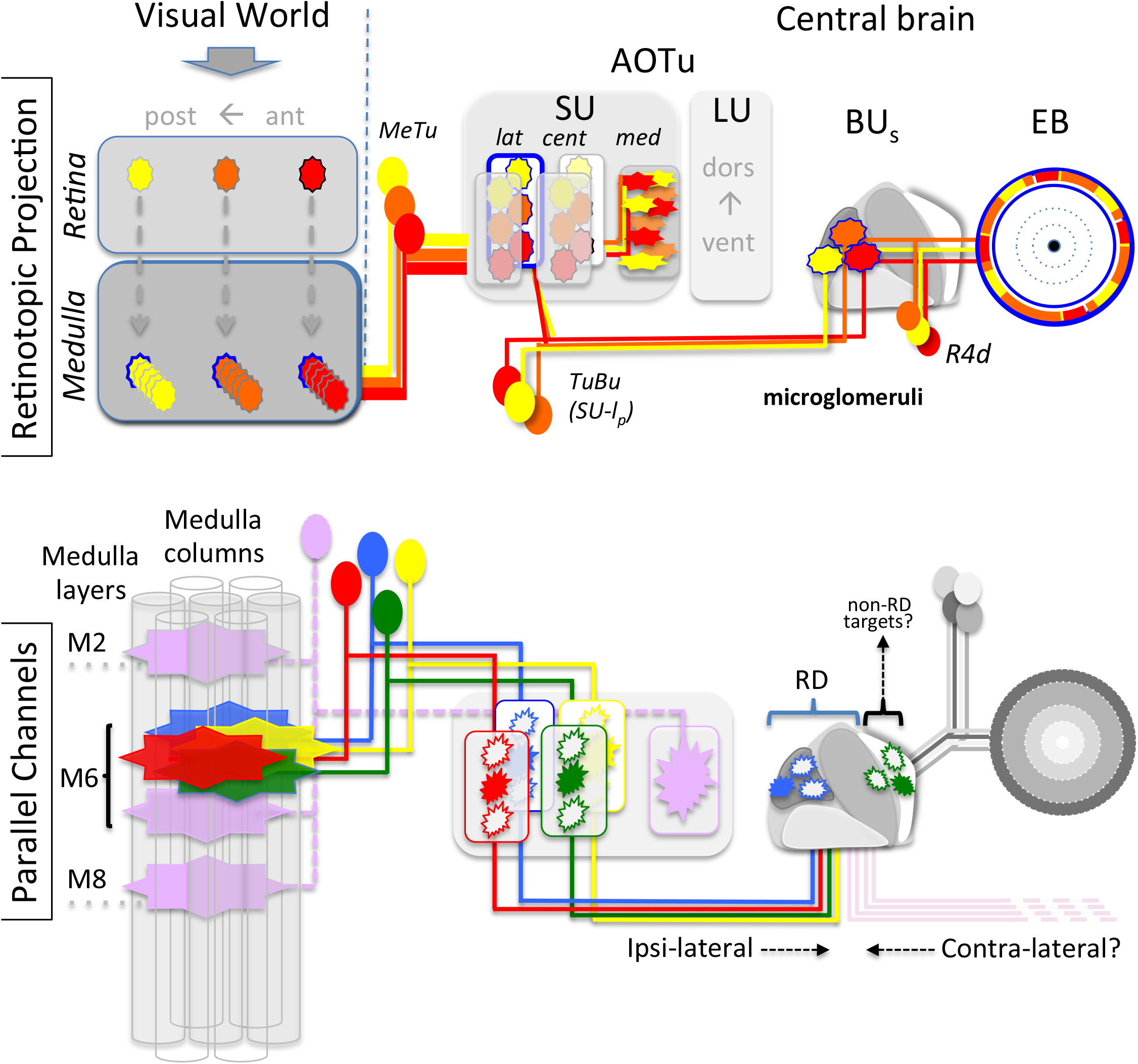

First, retinotopic representation in the dorsal half of the medulla is maintained in a defined sub-compartment of AOTu, spatially separated from lobula representation within the AOTu. Single cell analysis showed a strict topographic correlation between the anterior-posterior position of the dendritic fields of MeTu projection neurons in the medulla and their restricted axon termination along the dorso-ventral axis within the SU domains of the AOTu. Retinotopy is further conserved in the AOTu output neuron projections towards the bulb neuropil, where TuBu neurons synapse onto defined micro-glomeruli with dendrites of specific ring neurons. All ring neurons of the same type innervate a common ring layer within the ellipsoid body, raising the question as to what degree topographic information is integrated within central complex neuropils. One could speculate that when the morphology of axon terminal does not contain topographic features, spatial information might be translated into a defined sequence of alternating temporal activity of ring neurons.

Second, peripheral points in space are represented in multiple parallel channels within the AOTu domains but diverge within the bulb regions. Here we could identify not only a complete mapping of one topographic channel onto a defined ring neuron field (SU-l_post_ > R4d) but also a merely partial innervation of another ring domain by a second topographic MeTu channel (SU-c_ant_ > R2). In addition to the point-to-point representation via multiple topographic channels, the AOTu also contains a non topographic input, in which multi-columnar medullar projection neurons project onto a separate domain, which is further relayed to the non-topographic inferior bulb area. Interestingly, while other MeTu neurons (type-1 morphology) establish dendritic fields within a single medulla layer, the non-topographic channel is formed by MeTu-m neurons innervating the medial SU, which integrate three different medulla layers (type-2 morphology), reminiscent of lobular LC neurons. In fact, LC neurons are the main afferents of the adjacent AOTu large unit, for which a rough topographic pattern of along the dorso-ventral axis has been described (Wu, Nern, W Ryan Williamson, et al. 2016). In addition, MeTu-m neurons also develop a collateral arborization in the lobula, indicating that this pathway directly integrates visual information from the medulla and lobula. Furthermore, a small fraction of MeTu-m neurons also show restricted termination within the AOTu domain, indicating a more heterogeneous neuronal population. This generates an interesting sequence of adjacent domains within the AOTu, from exclusive topographic medulla input in SU-l and SU-c domains, a mixed zone of combined medulla/lobula integration within the SU-m domain, and a zone of exclusive lobula representation within the LU (Fig. 7). Thus, we could identify a unique visual circuitry with multiple parallel topographic pathways separated from nontopographic connections.

Based on their molecular identity, three principal types of medullar columnar neurons (MeTu) provide input into the AOTu, with overlapping dendritic fields within the medulla but segregated axon terminals to distinct AOTu domains. Type-1 MeTu classes, MeTu-l and - c, have a similar neuronal morphology with dendrite arborization only in a single medulla layer (M6) and restricted axon terminals in the AOTu domain, thereby building the multiple parallel topographic pathways (Fig. 7). In contrast, the MeTu-m class of medulla projection neurons (type-2) integrates multiple medulla layers and arborizes broadly in the AOTu domain. As the organization of the MeTu-l and MeTu–c neurons defines a point-to-point projection similar to the columnar identity in the medulla, these AOTu domains are constituted to relay topographic information towards the central brain. Interestingly, the axons of single MeTu-m neurons converge and intermingle within their AOTu domain, which is reminiscent of the afferent organization of LC neurons from the lobula within optic glomeruli in the PVLP region. In fact, LC neurons are the main afferents of the adjacent AOTu large unit (LU), for which a rough topographic pattern along the dorso-ventral axis has been described (Wu, Nern, W Ryan Williamson, et al. 2016). In addition, MeTu-m neurons also develop a collateral arborization in the lobula, indicating that this pathway directly integrates visual information from the medulla and lobula. Furthermore, a fraction of MeTu-m neurons also shows restricted termination within the AOTu domain, indicating a more heterogeneous neuronal population. This generates an interesting sequence of adjacent domains within the AOTu, from exclusive topographic medulla input in SU-l and SU-c, a mixed zone of combined medulla/lobula integration within SU-m, and a zone of exclusive lobula representation within the LU (Fig. 7).

The borders of the SU compartments are well respected also by the projecting TuBu neurons (Fig. 4C-D), an observation that takes the concept of parallel pathways to the next synaptic step, the bulb. While this neuropil with its afferent (TuBu) and efferent (R neurons) channels has been intensively studied recently (Seelig & Jayaraman 2013; Seelig & Jayaraman 2015; Omoto et al. 2017; Sun et al. 2017), there is still lack of knowledge of the precise connections formed, that translate into the central complex. Four major findings of the TuBu-R neuron circuit are provided by our study: First, the topographic position of TuBu dendrites in the SU is not translated into a defined position within the bulb, but exhibits a targeting plasticity within a restricted bulb area (Fig. 5). Second, with exception of the medial domain, all TuBu classes we analyzed project into the BUs, but into defined regions without intermingling (Fig. 4 and data not shown). Third, TuBu classes project onto dendritic areas of R neuron classes (‘sectors’) within the bulb, and we describe specific connections formed between TuBu neurons and R neuron classes (Fig. 6). Fourth, although we could associate three R neuron classes with the BUs, a considerable amount of TuBu neurons (TuBu-c_a_) is targeting a medial bulb region with unknown postsynaptic partners, suggesting non-R neurons are also contacted directly by TuBu neurons (Fig. 6J-L). Furthermore, as R2 covers a large area of the bulb, it is likely that several TuBu-classes are integrated in one ring structure. In addition, there are contralateral neurons described in the locust (el Jundi & Homberg 2012) and bumblebee (Pfeiffer & Kinoshita 2012), which connect the AOTu units of both hemispheres. As contralateral inhibition is a feature of bulb neurons (Sun et al. 2017) and there are large, uncovered sectors in the BU_s_, we speculate that also projections of contralateral neurons are part of the bulb network.

While the recent demonstration of AOTu-EB pathways with the bulb as a tripartite structure (Omoto et al. 2017) included both afferent and efferent neurons, we would like to refine this picture by highlighting that, although our analysis of TuBu-neurons is mainly constricted to two TuBu classes (one in the SU-l_post_ and the other in the SU-c_ant_ domain), both neural ensembles target areas within the superior bulb (BUs). Broader expression lines also showed exclusive TuBu neuron innervation of the BUs, indicating further TuBu classes targeting this bulb area (data not shown). Thus, we expect at least four different classes of TuBu neurons to exclusively innervate the BUs, connecting each to a different set of output neurons, indicating a more intricate organization of the bulb, in particular the BUs. Furthermore, as in our screen we did not encounter TuBu neurons targeting the BUa (compare Omoto et al. 2017), this holds the possibility that the AOTu-SU consists of even further functional units. Future studies on activation patterns of the distinct TuBu neuron classes are expected to shed light on which stimuli are percieved by the distinct pathways. While functional differences between the BUi and BUs have been described (Omoto et al. 2017), their functional studies and also those of other groups (Seelig & Jayaraman 2013; Sun et al. 2017) do not resolve neuronal activity of TuBu classes and the responses of R neurons within the BU_s_. We favor a model where retinotopic information in the BUs remains represented in the respective sectors, associated with their TuBu class.

The principle pathway, involving AOTu as a central relay station connecting medullar/lobular input with central neuropils, is widely shared among different insect taxa, where homologous structures can be found, e.g. orthopterans (Homberg et al. 2003), hymenopterans (Mota et al. 2011) and beetles (Immonen et al. 2017). Few insect species have so far been analyzed in terms of stimuli conveyed by this pathway, however, different compartments are associated with celestial orientation using polarized light (Pfeiffer et al. 2005) or in chromatic processing (Paulk et al. 2008; Mota et al. 2013). Dorsal rim neurons sensitive for polarized light are crucial for the sky-compass orientation conveyed by the AOTu pathway in most insects analyzed, like locusts (Pfeiffer et al. 2005), butterflies (Heinze & Reppert 2011) and honeybees (Held et al. 2016), but this task is likely not fulfilled by the AOTu in Drosophila (Omoto et al. 2017). In addition, the neural circuit upstream of MeTu neurons remains unknown. While chromatic processing is accomplished in other insects via the AOTu pathway, we did not find support for a direct interaction of the color-vision pathway in the medulla layer M6 (Fig. 2L). The subdivision as described above is a common feature of the insect AOTu. We present here a combination of cell-surface molecules that can serve as landmarks for the AOTu-subdivision and further refinement by genetic labeling revealed ‘territories’ of distinct neuronal classes in a constricted volume of a neuropil. In addition, as the fruit fly is among the smallest species for which the AOTu has been analyzed, a more sophisticated architecture of the SU-homologue might be revealed in other insect taxa. In addition, we describe a distinct expression pattern of cell-surface molecules among the entirety of optic glomeruli (Fig. 1E and data not shown). The molecular mechanism that underlie the formation of the LC-optic glomeruli network remain to be resolved. On the anatomical and functional level, optic glomeruli share many features with the synaptic neuropil within the antennal lobe, which led to the postulation that the glomerular organization in the protocerebrum (optic glomeruli) and the deutocerebrum (olfactory glomeruli) are homologous structures (Strausfeld 1989; Mu et al. 2012). We found molecular characteristics in the PVLP and AOTu that resemble the combinatorial code of cell-surface proteins in the olfactory system (e.g. expression patterns of Ten-m, Con, Caps and Sema1a in both systems). First developmental studies of mutant LC and MeTu neurons (unpubl.) suggest common mechanisms of glomerular targeting in both sensory systems. Although the idea of a serial homolgy of glomerular organized neural system is far from being resolved, it will be intriguing for further studies to analyze the developmental mechanisms that underlie the circuit formation of these parallel AOTu pathways and optic glomeruli circuits as well as to compare them with known molecular functions during olfactory system maturation.

## MATERIALS AND METHODS

### Fly rearing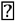

Flies were maintained in vials containing standard fly food medium at 25°C unless otherwise mentioned. Canton-S flies were used as a wild type strain.

### Fly stocks

The following driver lines were generated at the Fly Light Gal4-/LexA-Collection (Jenett et al. 2012) and obtained from Bloomington Drosophila Stock Center (BDSC). One driver line was obtained from the Vienna Tiles collection (Kvon et al. 2014).

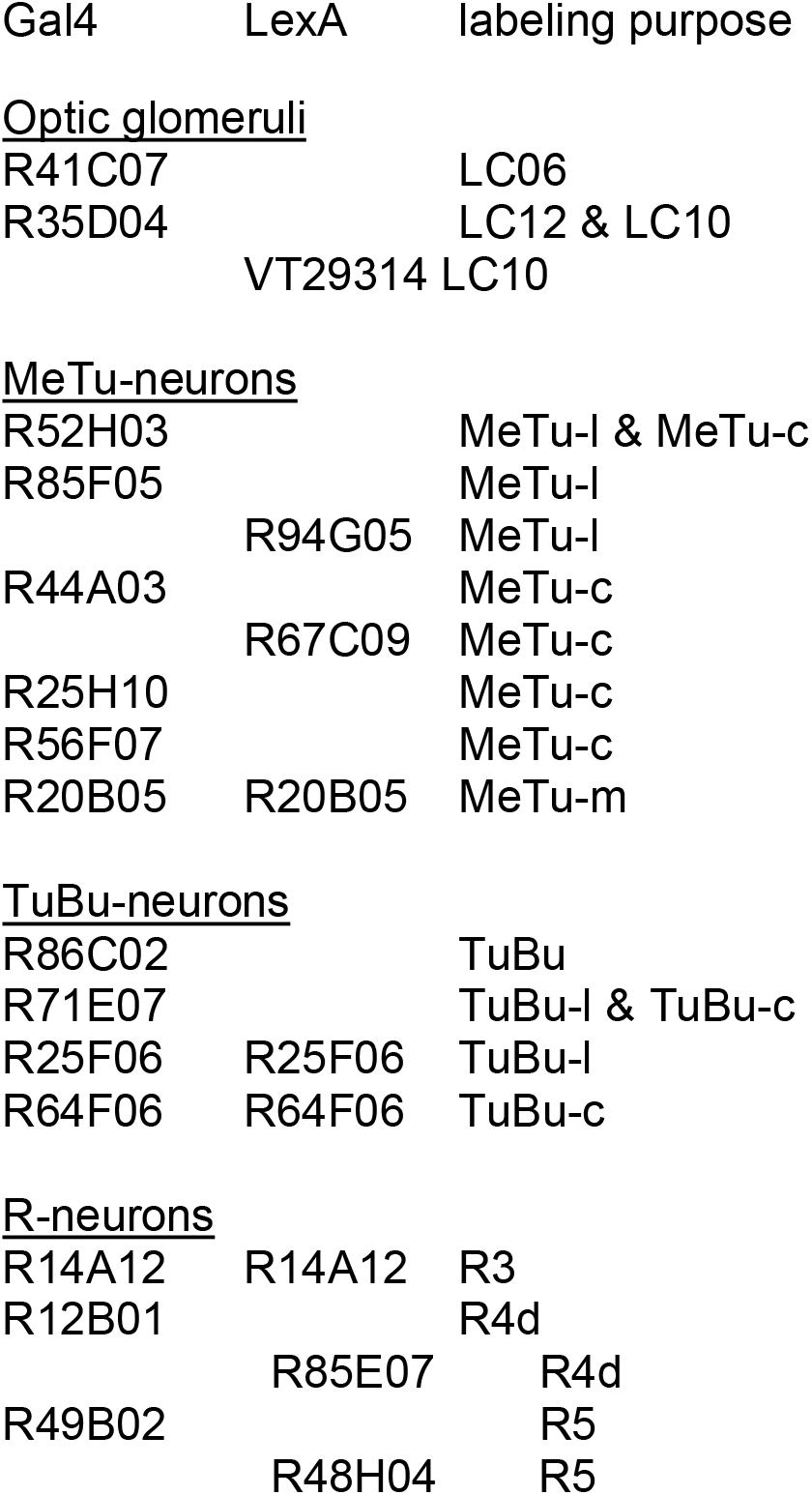

| Gal4 | LexA | labeling purpose |
| --- | --- | --- |
| Optic glomeruli |  |  |
| R41C07 |  | LC06 |
| R35D04 |  | LC12 & LC10 |
|  | VT29314 | LC10 |
| MeTu-neurons |  |  |
| R52H03 |  | MeTu-l & MeTu-c |
| R85F05 |  | MeTu-l |
|  | R94G05 | MeTu-l |
| R44A03 |  | MeTu-c |
|  | R67C09 | MeTu-c |
| R25H10 |  | MeTu-c |
| R56F07 |  | MeTu-c |
| R20B05 | R20B05 | MeTu-m |
| TuBu-neurons |  |  |
| R86C02 |  | TuBu |
| R71E07 |  | TuBu-l & TuBu-c |
| R25F06 | R25F06 | TuBu-l |
| R64F06 | R64F06 | TuBu-c |
| R-neurons |  |  |
| R14A12 | R14A12 | R3 |
| R12B01 |  | R4d |
|  | R85E07 | R4d |
| R49B02 |  | R5 |
|  | R48H04 | R5 |

#### Stocks for clonal analysis and effector lines for cell labeling

*FRT42D; FRT42D TubP-Gal80; UAS-mCD8::GFP, UAS-mCherry* were obtained from BDSC. The *UAS*-DenMark construct was kindly provided by Bassem Hassan, *LexAop::GFP* was a kind gift from Andrew Straw. Flies for synaptic-GRASP experiments (UAS-Syb::spGFP1-10 & LexAop spGFP_11_::CD4) (Karuppudurai et al. 2014) were a kind gift from Chi-Hon Lee.

#### Specific cell labeling

*Or67d::GFP and OR67d-Gal4* (Couto et al. 2005) was used for olfactory class visualization, glia cells were marked by repo-*Gal4* (Sepp et al. 2001) and Chi-Hon Lee provided the *ort^C1a^-LexA::VP16* (Ting et al. 2014) construct for labeling of Dm8 neurons.

Caps-Gal4 (Shinza-Kameda et al, 2006) ???

### Antibodies used in this study

Primary antibodies used were: 24B10/Mouse anti-Chaoptin (1:50, DSHB), DN-Ex #8/Rat anti-CadN (1:20, DSHB), Flamingo#74/Mouse anti-flamingo (1:20, DSHB), Rabbit anti-GFP (1:1000, Invitrogen, Carlsbad, CA). Mouse anti-Teneurin-m (1:20) was a kind gift from Stefan Baumgartner, anti-Connectin (1:20) was kindly provided by Robert AH White.

Following secondary antibodies were used: Goat anti-Rabbit Alexa-488 (1:500), Goat anti-Rabbit Alexa-568 (1:300), Goat anti-Mouse Alexa-488 (1:300), Goat anti-Mouse Alexa-647 (1:500), Goat anti-Rat Alexa-647 (1:300). All secondary antibodies were obtained from ThermoFisher Scientific (Alexa Fluor®, Molecular Probes™).

### Clonal analysis

Two approaches for visualization of large and small genetic mosaics were used respectively. For labeling larger mosaics, MARCM clones with an ey-Flp insertion on the X chromosome (Newsome et al. 2000) were generated as previously described (Lee & Luo 1999). For small clones and single-cell analysis, we used the temperature-sensitive hs-mFlp5 promotor in combination with a FLYBOW (FB1.1B)-construct (Hadjieconomou et al. 2011; Shimosako et al. 2014). Prior to screening for brains with single cell labeling, a heat shock was given to developing flies (L2-stage, L3-stage, early pupal) for 30min, 1h or 2h at 38°C. The pupae were then allowed to further develop at 25°C and dissected within two days after eclosion.

### Immunohistochemistry

Drosophila brains were dissected in phosphate-buffered saline (PBS) and fixed in 4% paraformaldehyde (PFA) in PBS for 20 min. Samples were washed 3 × 15 min with PBS-T (PBS containing 0.3 % Triton X-100) and blocked for 3 hours (10% Goat serum in PBS-T) under constant shaking on a horizontal shaker, before incubating in primary antibody solution for two days at 4°C. Washing procedure was repeated before incubating with secondary antibody for two days at 4°C. Following three times washing, the brains were mounted in Vectashield® (Vector Laboratories, Burlingame, CA) anti-fade mounting medium prior to confocal microscopy. Images were obtained using a TCS SP5II confocal microscope (Leica) using 20x and 63x glycerol immersion objectives. Image processing was performed using ImageJ and Adobe Photoshop® CS6.

